# Venom gland organogenesis in the common house spider

**DOI:** 10.1101/2024.03.14.585096

**Authors:** Afrah Hassan, Grace Blakeley, Alistair P. McGregor, Giulia Zancolli

**Affiliations:** Department of Ecology and Evolution, University of Lausanne, 1015, Lausanne, Switzerland; Oxford Brookes University, Oxford, United Kingdom; Department of Biosciences, Durham University, South Road, Durham, DH1 3LE, United Kingdom; Swiss Institute of Bioinformatics, 1015, Lausanne, Switzerland

**Keywords:** Developmental origins, evolutionary innovation, HCR, toxins, spiders

## Abstract

Venom is a remarkable innovation found across the animal kingdom, yet the evolutionary origins of venom systems in various groups, including spiders, remain enigmatic. Here, we investigated the organogenesis of the venom apparatus in the common house spider, *Parasteatoda tepidariorum*. The venom apparatus consists of a pair of secretory glands, each connected to an opening at the fang tip by a duct that runs through the chelicerae. We performed bulk RNA-seq to identify venom gland-specific markers and assayed their expression using RNA *in situ* hybridisation experiments on whole-mount time-series. These revealed that the gland primordium emerges during embryonic stage 13 at the chelicera tip, progresses proximally by the end of embryonic development and extends into the prosoma post-eclosion. The initiation of expression of an important toxin component in late postembryos marks the activation of venom-secreting cells. Our selected markers also exhibited distinct expression patterns in adult venom glands: *sage* and the toxin marker were expressed in the secretory epithelium, *forkhead* and *sum-1* in the surrounding muscle layer, while *Distal-less* was predominantly expressed at the gland extremities. Our study provides the first comprehensive analysis of venom gland morphogenesis in spiders, offering key insights into their evolution and development.

## Introduction

Across the tree of life, many organisms have independently acquired the capability to produce and deliver cocktails of bioactive molecules, either for predation or defence purposes, making venom one of the most common secretions in nature^1^. In most venomous animals, venom is synthesised in specialised exocrine glands and subsequently administered into other organisms via dedicated delivery structures such as fangs and stingers^1^.

Venoms are primarily studied for their pharmacological applications, biodiscovery and drug development^2–4^, although they also play an important role in fundamental research, serving as models for understanding molecular evolution and ecological dynamics^5,6^. Despite their significance, the evolutionary and developmental origins of many venom systems remain poorly understood^6^.

Among venomous animals, spiders have emerged as excellent experimental systems for investigating developmental mechanisms from an evolutionary standpoint. Notably, studies of the central American wandering spider *Cupiennius salei* and the common house spider or cobweb spider *Parasteatoda tepidariorum* have significantly contributed to the field of evolutionary and developmental biology (‘Evo-Devo’)^7–9^. Innovations such as silk and venom have enabled spiders to become among the most successful predators on land^10–13^ and they therefore represent excellent models for studying the origins of evolutionary novelties. However, the evolutionary and developmental origins of such key evolutionary innovations, particularly the venom system, remains largely unexplored.

In spiders, the venom apparatus is associated with the chelicerae, the first pair of appendages, commonly referred to as jaws^14^ (Fig. 1). The chelicerae feature a pair of fangs which are used to deliver venom through an opening located at their tips. Venom is produced in the venom glands, a pair of exocrine glands, each connected to the fang’s opening by a duct that runs through the chelicerae^14^ (Fig. 1b). In Mesothelae, a sub-order of spiders characterised by ancestral features and branching basally from other extant taxa, the venom glands are located at the base of the chelicerae^15^. In Mygalomorph spiders, including tarantulas, funnel web spiders, and trapdoor spiders, the venom glands extend throughout the chelicerae. In contrast, Araneomorphs, which are considered more derived and represent most of the living species, have venom glands that extend further into the prosoma^14^. For a schematic representation, see Figs 2 and 3 in Lüddecke et al^13^.

**Figure 1.**
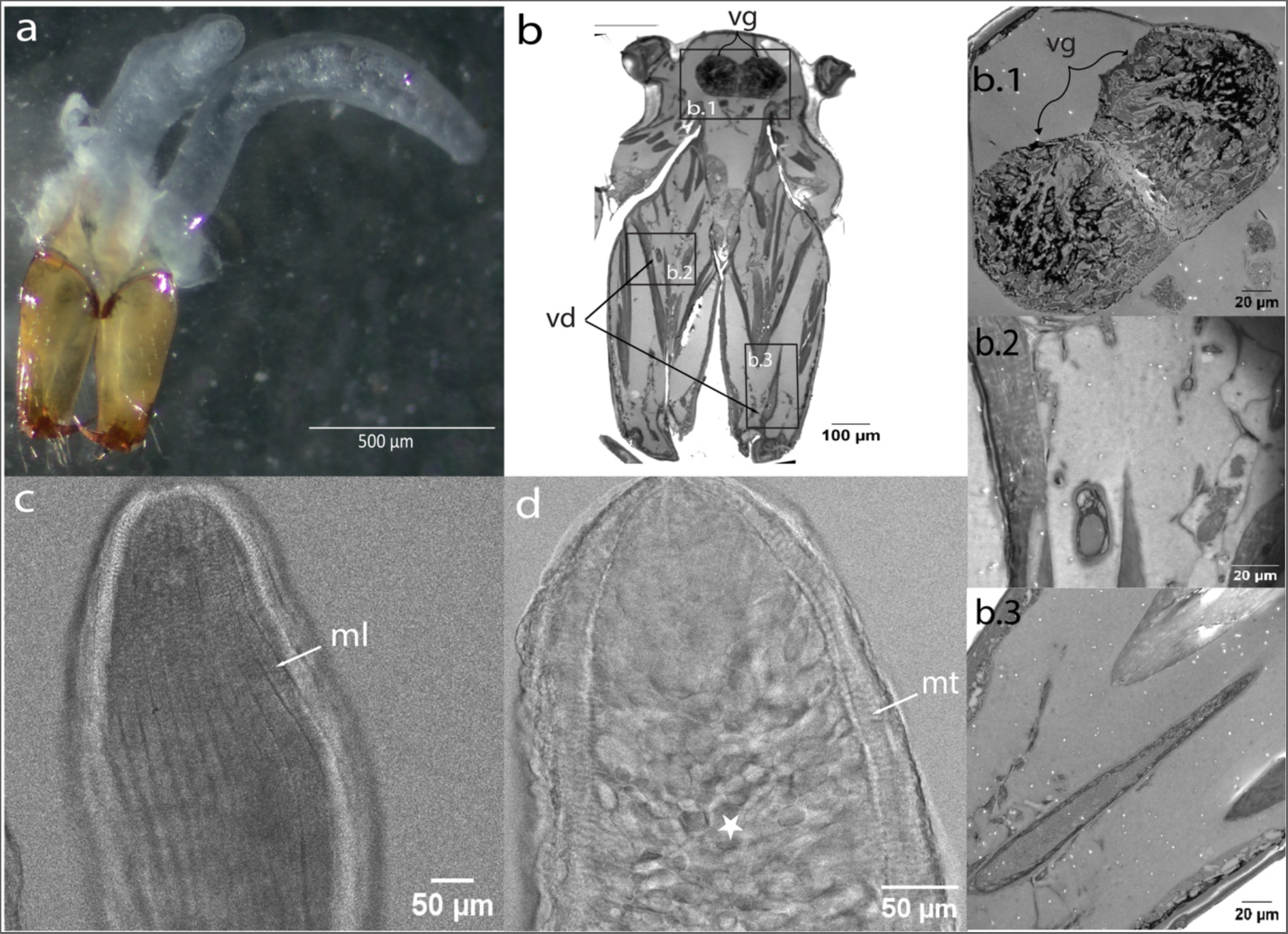
Venom apparatus of adult *P. tepidariorum*. a) Overview of the whole venom apparatus, with a pair of elongated, tubular glands above the chelicerae. b) Histological frontal section across a spider head showing the anterior most region of the venom glands (vg) in the dorsal side of the prosoma, and the venom duct (vd) within the chelicerae. Higher magnification images of the venom glands (b.1), and the venom ducts (b.2, b.3). c) Confocal LSM scan revealing an outer layer of longitudinal muscle (ml) tightly surrounding the venom gland. d) Underneath the longitudinal muscle layer, transversal muscle fibres (mt) and connective tissue surround the secretory tissue characterised by an alveolar structure (star).

**Figure 2.**
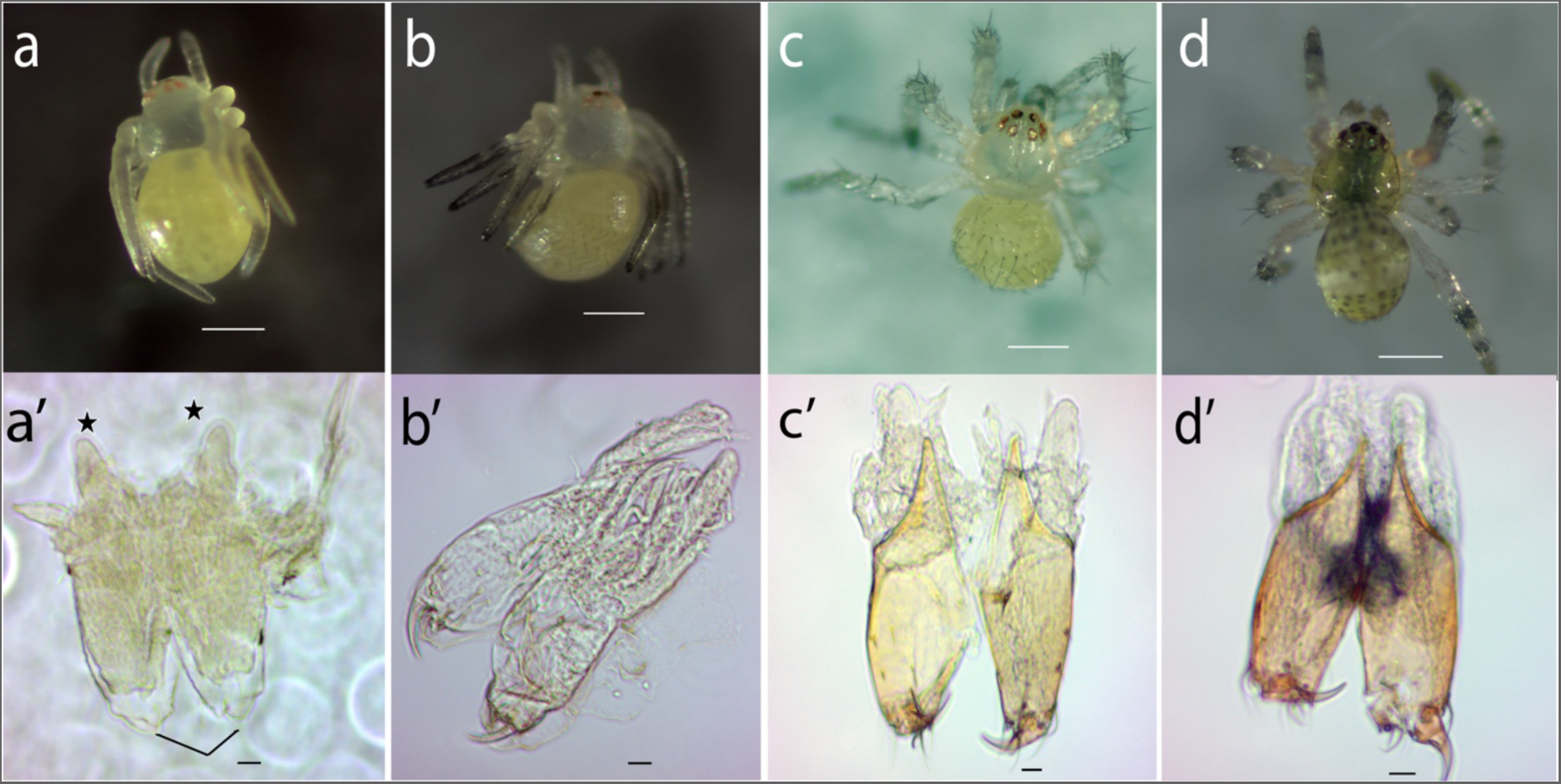
Postembryos and first instars of *P. tepidariorum* and their venom apparatus. a) Early postembryo (post-hatching) and a’) its chelicerae. Most early postembryos lie upside down, immobile, though their legs exhibit some twitching movements. The developing fangs (arrows) and venom glands (stars) already outside of the chelicerae are indicated. b) Late postembryo (1.5 days) with sensory hairs on the opisthosoma and legs. The hairs primarily appear on the posterior side of the opisthosoma and legs, which become darker, particularly at the extremities. Additional hairs are visible on the chelicerae beneath the postembryonic membrane. Legs movements become more frequent, although their joints are not completely developed. b’) Sharper, more defined fangs, and sensory hairs at the tip of the chelicerae. c) Early first instar (3 days) characterised by more developed legs. The body remains unpigmented, although the opisthosoma starts to exhibit a yellowish hue, and the legs become thinner and longer. c’) Pigmented chelicerae and elongated venom glands. d) Late first instar (5 days) with a pigmented prosoma. The opisthosoma develops dark blotches, with a patch appearing in the centre of the prosoma. d’) Fully developed venom apparatus. a-d scale bar: 200 *μm*; a’ to d’ scale bar: 20 *μm*.

**Figure 3.**
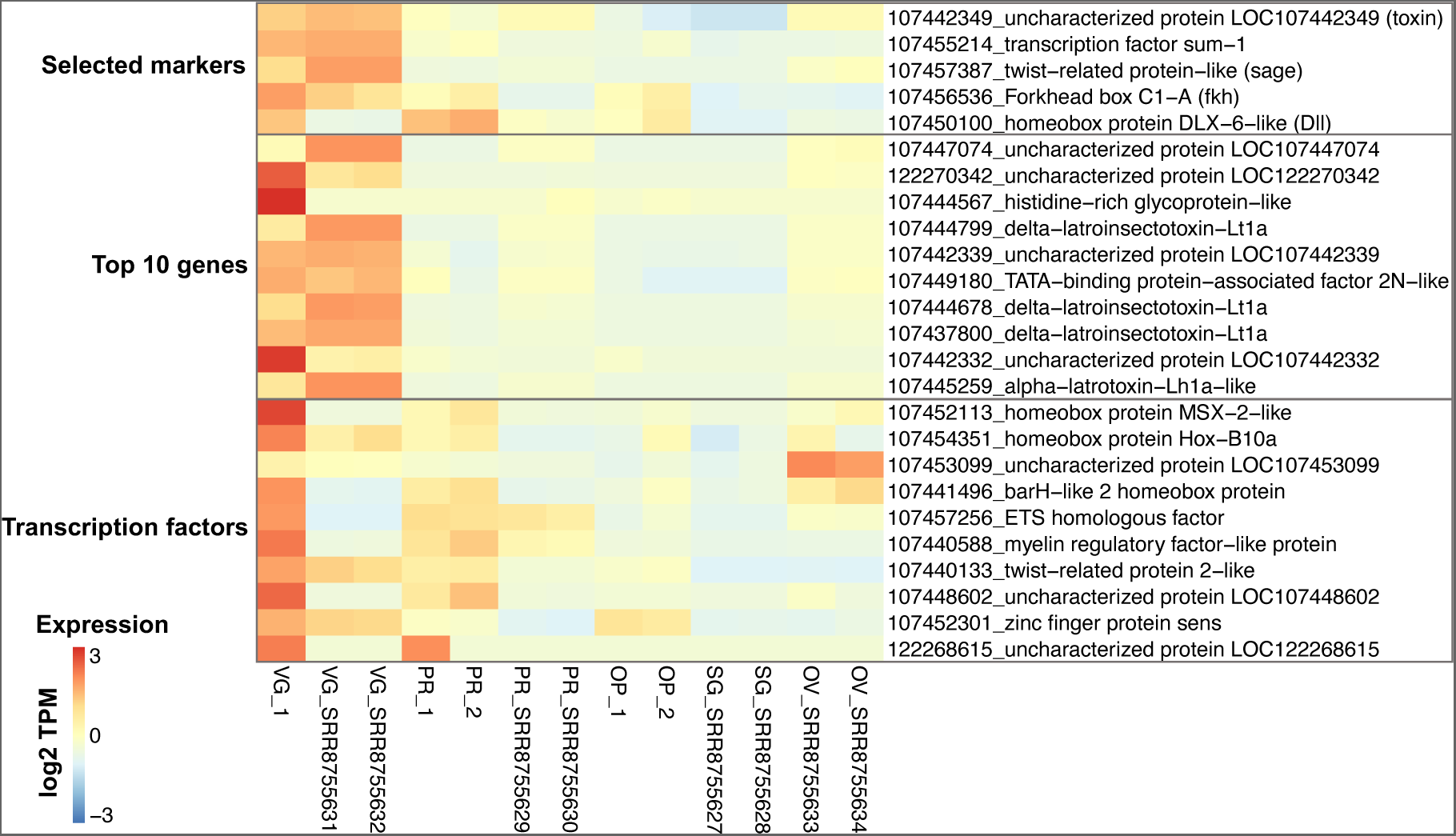
Heatmap of venom gland specific genes. The top five genes correspond to the markers selected for the *in situ* HCR experiments. In brackets are the names used to refer to the genes in the text. The genes below represent the top ten venom gland-specific genes (highest FC) in *P. tepidariorum* juveniles. The last set of genes correspond to the juvenile venom gland-specific transcription factors (FC and tpm >= 2). Samples numbered “_1” and “_2” are the juveniles from this study. VG = venom glands; PR = prosoma; OP = opisthosoma; SG = silk glands; OV = ovaries. The adults are from SRA libraries generated in ^21^. Expression values are row scaled log2(TPM+1).

Rempel observed that in *Latrodectus mactans* “invaginations in the chelicerae disclose the origin of the poison glands” in 370 hour-old embryos^16^. In two species of *Peucetia*, rudimentary venom glands were observed in first instar juveniles in the form of a short tube slightly enlarged at the distal end of the chelicera, although it did not reach the fang^17^. Venom glands were found in the early first pre-larval stage of *Phoneutria nigriventer*, with the entire venom apparatus becoming fully formed in the stage preceding eclosion from the cocoon^18^. Despite these observations, there has not been a systematic investigation of the time-series of the development of the spider venom apparatus. Moreover, these observations do not conclusively clarify whether the venom gland primordium originates distally or proximally in the chelicera.

In addition to the uncertainties surrounding the organogenesis of spider venom glands, their evolutionary origins also remain enigmatic^6^. The current debate revolves around whether the ancestral organs of origin were salivary glands or silk glands^13^, although a recent study found support for the silk gland origin hypothesis based on similarities in the transcriptomes of these two organs^19^.

In this study, we aimed to bridge this knowledge gap by providing a more comprehensive description of the development of the venom apparatus in *P. tepidariorum* using both morphological observations and analyses of expression patterns of selected markers at different time points during embryonic and post-embryonic development stages as well as in adult venom glands.

## Results

### Morphology of the developing venom apparatus in *P. tepidariorum*

To better understand the organisation of the venom apparatus in *P. tepidariorum* and trace its developmental origin, we initially examined the chelicerae and venom glands in postembryos, corresponding to the stage following eclosion, first instars and adults^20^. The postembryos and first instar stages were further categorised into ‘early’ and ‘late’ stages based on specific developmental features (see below).

Eggs typically hatch 7 to 8 days after the cocoon is laid^20^. Immediately after eclosion, early soft-bodied postembryos appear with a transparent prosoma, a milky opisthosoma, and red eyes (Fig. 2a). At this stage, the chelicerae are transparent, and the fangs are soft and rounded rather than sharp, remaining concealed underneath the postembryonic membrane. Short venom glands can be observed atop the chelicerae below the eyes (Fig. 2a’).

Around 1.5 days post-hatching, the appearance of sensory hairs marks the transition to the late postembryonic stage (Fig. 2b). While still unpigmented and partially covered by the membrane, the fangs are now more defined, slightly harder and sharper compared to the earlier stage (Fig. 2b’).

After about 3 days post-hatching, the spiders emerge from the cocoon and begin producing silk threads that they use to position themselves upside down. These early first instars possess more fully developed legs capable of supporting the spider’s weight (Fig. 2c). The chelicerae darken as the cuticle begins to harden, and the fangs start to take on a brown coloration. Sensory hair appears at the base of the fangs. The venom glands become more elongated compared to postembryos and begin to extend into the prosoma (Fig. 2c’).

The transition to late first instars, at around 5 days post-hatching, is marked by body pigmentation, although the legs remain transparent (Fig. 2d). The chelicerae continue to harden, and the fangs are darker. The venom glands become more conspicuous in volume, approximately matching the length of the chelicerae, and continue to extend into the prosoma (Fig. 2d’).

In adults, the venom glands are elongated and occupy the upper part of the prosoma along its entire length (Fig. 1 a,b). Confocal laser scanning microscopy (CLSM) of dissected whole-mount venom glands revealed an outer layer of longitudinal muscle fibres (Fig. 1c) and an inner layer of transversal muscle fibres tightly surrounding the glandular tissue, which is organised into secretory units (Fig. 1d).

### Gene expression analysis by bulk RNA-seq

To identify developing venom gland-specific markers, we dissected spiders at four days post-hatching (late first instar stage) and extracted mRNA from the venom glands, the rest of the prosoma, and the opisthosoma. Due to the minute size and delicate nature of the venom glands, they were left attached to the chelicerae. We obtained two replicates for the prosoma and opisthosoma, but only one sample for the venom gland. The RNA-seq libraries ranged in size from 45 M to 61 M reads.

In total, 11,155 genes with Transcript Per Million (TPM) >= 2 were expressed in the juvenile venom gland, which was comparable to the other samples (11,934 for the prosoma and 10,162 for the opisthosoma) (Supplementary Dataset S1,S2). To identify tissue-specific genes, we calculated the difference in expression between the venom gland sample and the tissue with the second-highest expression value. Genes with fold change (FC) and TPM >= 2 were considered as tissue specific, and those with the highest values were listed as candidate venom gland markers (Supplementary Dataset S3). In total, we found 426 venom gland-specific genes.

Since we had a single replicate for the venom gland, we compared our gene expression data from juveniles to eight published libraries from adult specimens to help validate the specificity of venom gland markers. These libraries included samples from venom glands, prosoma, ovaries and silk glands^21^, and we mapped them to the Ptep_3.0 transcriptome (GCF_000365465.3) in the same way as we did for our samples. Overall, juvenile and adult venom glands exhibited similar expression patterns, with 84% of the genes expressed in the adult also expressed in the juvenile. However, fewer genes were expressed in the adult gland (6,208), possibly because of more precise dissections^21^. Similarly, we found fewer venom gland-specific genes (n = 239) in the adult, and of these, 94 were also specific in the juvenile. Several transcription factors were specific in the juvenile but not in the adult gland, including some homeobox genes such as *Msx-2*, *Hox-B10a* and *BarH-2*, possibly due to contamination of mRNA from the chelicerae (Fig. 3). Among the genes with the highest TPM values in juvenile venom glands were several annotated as ‘uncharacterised proteins’, many of which have been found secreted in the venom (e.g., LOC107442349)^21^. Among the genes annotated as toxins (e.g., latrototoxin, latrocrustotoxin), we found 101 which were expressed in the juvenile venom gland; of these, 19 were gland-specific (FC > 2). In the adult, we found 44 toxin genes, 30 of which were also expressed in the juvenile venom gland. Many of the uncharacterised genes, as well as ‘TATA-binding protein-associated factor 2N-like’, which were specific in both juvenile and adult glands, lacked orthologs in other species according to the OrthoDB annotation.

Considering the uncertainty regarding the function of genes annotated as either uncharacterised or lacking orthology, we opted to focus solely on the expression of a few known genes, specifically transcription factors, considering their crucial role in organ development and cell differentiation. To identify transcription factors, we initially focused on genes listed in the KEGG orthology KO 03000 ‘Transcription factors’. Among these, twelve were upregulated in juvenile venom glands (Fig. 3). *sum-1* had the highest FC, and its specificity to venom glands has been observed in other spider species^22^. *forkhead box C1-A* displayed a similar pattern, with venom gland-specific expression in both juveniles and adults (Fig. 3). This gene is ortholog to the *Drosophila melanogaster forkhead* (*fkh*) (and so we use this name hereafter), which works with the product of *sage* to activate the expression of salivary gland-specific enzymes and secreted proteins^23^. The ortholog of *sage* in *P. tepidariorum* is a gene named ‘twist-related protein-like’, which was not categorised as a transcription factor in the KEGG database. However, considering the important regulatory roles of *fkh* and *sage* in *Drosophila* salivary gland determination, and the upregulation of *fkh* in venom glands, we investigated the expression pattern of twist-related protein-like (referred to as *sage* from here onwards) and found that it was highly upregulated in both juvenile and adult venom glands (Fig. 3). Consequently, we selected *sum-1*, *fkh* and *sage* as candidate venom gland markers for *in situ* hybridisation chain reaction (HCR) experiments, as well as one toxin gene (LOC107442349) to elucidate when the epithelial secretory cells begin producing venom. In addition to the venom gland markers identified by RNA-seq analysis, we investigated the expression pattern of *Distal-less* (*Dll*) since this gene specifies the proximal-distal axis and is expressed in the embryonic appendages including the chelicerae^24–26^.

### Spatiotemporal expression of marker genes using *in situ* HCR

The experiments were divided into two multi-HCR sets: set 1 comprising toxin, *sum-1* and *Dll*, while set 2 included *sage* and *fkh*.

#### Adult venom glands

We started by assessing the expression patterns of the selected markers in dissected adult venom glands (Fig. 4, Supplementary Fig. S1). The toxin gene exhibited expression across the entire length of the gland, albeit non-uniformly, with a predominant signal in the secretory cells surrounding the lumen, particularly at both extremities of the gland (Fig. 4a). *sum-1* displayed uniform expression within the muscular layer surrounding the glandular epithelium (Fig. 4a). *Dll* expression was overall weak, but we detected strong, localised signals at both extremities of the gland, although this was more prominent at the distal end. We did not detect any signal in the middle of the gland. Additionally, *Dll* co-localised with the toxin, suggesting it is also expressed in secretory cells (Fig. 4a).

**Figure 4.**
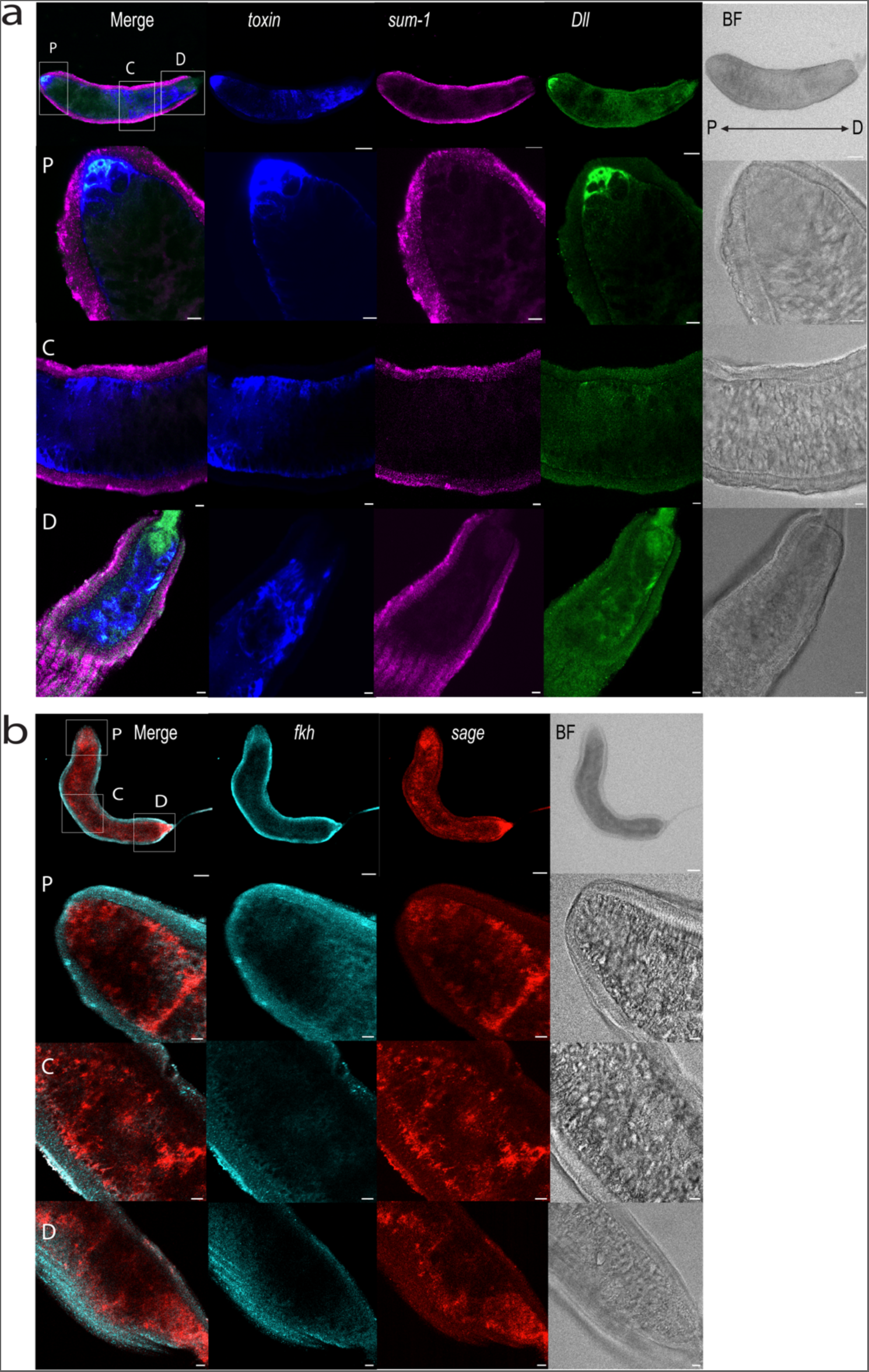
Expression of venom gland markers in adult venom glands. a) Whole-mount venom gland with *Dll*, *sum-1* and toxin HRC signals merged, the individual markers separately, and the bright field (BF) image. Scale bars: 100 *μM*. Rows below are 40X magnification of the distal (D), central (C), and proximal (P) regions. Scale bars: 20 *μM*. b) Whole-mount venom gland with *fkh* and *sage* expression merged, the individual markers separately, and the bright field (BF). Scale bars: 100 *μM.* Rows below are 40X magnification of the distal (D), central (C), and proximal (P) regions. Scale bars: 20 *μM*.

*sage* and *fkh* were expressed in distinct patterns in the adult glands (Fig. 4b). *fkh* gave low signal intensities and, akin to *sum-1*, was predominantly expressed within the muscular layer surrounding the glandular tissue. In contrast, *sage* was widely expressed within the secretory tissue.

#### Postembryos and first instars

After assaying the expression of the selected markers in adult venom glands, we proceeded with whole-mount HCR experiments on post-hatched juvenile spiders. We mounted the prosoma with either the dorsal or ventral side down, but we only detected signals when the spiders were mounted in the latter orientation.

In early postembryos, *sum-1* formed a ring-like expression domain, a similar expression pattern to that observed for this gene in adult glands within the muscular layer (Fig. 5a). No expression of the toxin marker or *Dll* was detected (Fig. 5a). In late postembryos, the toxin was expressed in the glands, and was surrounded by a ring of *sum-1* expression, consistent with previous observations in adults, with the toxin localised within the glandular tissue, and *sum-1* surrounding it (Fig. 5a). At this stage, *Dll* was expressed at the tip of the chelicerae. *sage* signal was notably strong in postembryos, especially at the early stage (Fig. 5b).

**Figure 5.**
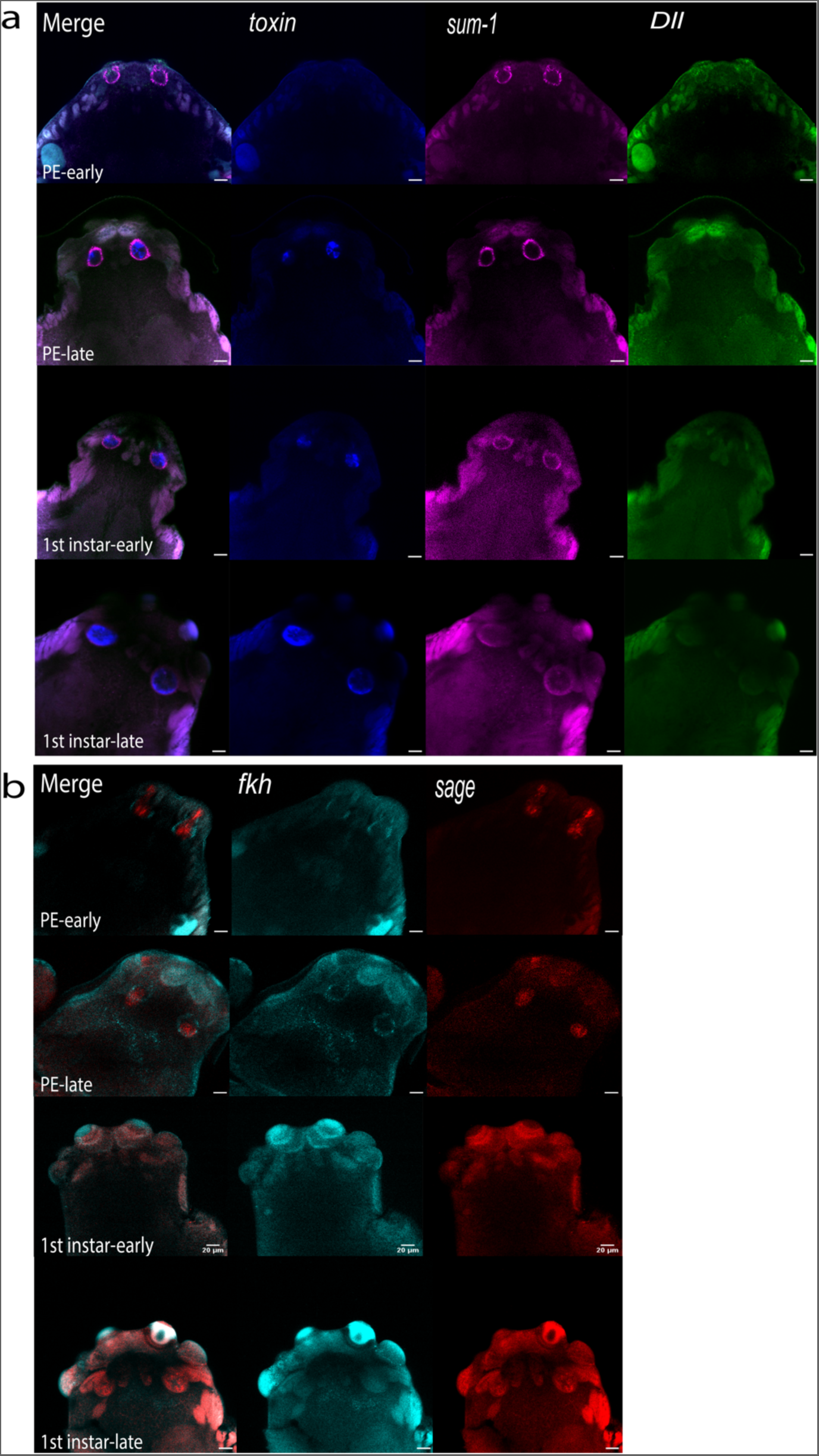
Expression of venom gland markers in postembryos and first instars. a) Expression of toxin, *sum-1* and *Dll* in first instars late and early stages, and postembryos (PE) late and early stages. b) Expression of *fkh* and *sage* in first instars late and early stages, and postembryos (PE) late and early stages. Scale bar 20 μm.

Consistent with observations in adult glands, *sage* signal was localised within the gland, while *fkh* exhibited expression in a ring around it, similar to *sum-1* (Fig. 5b). Additionally, we observed a signal at the sides of the prosoma, at the level of the coxae. However, it was difficult to distinguish this signal from background. At this stage, the glands had elongated into the prosoma because the signals were located outside the chelicerae and in the prosoma.

In both early and late first instars, *sum-1* exhibited the typical ring-like expression around the toxin signal in the prosoma (Fig. 6a). *Dll* expression was either absent or very weak and localised to the extremities of the gland as well as being expressed in the fangs (Supplementary Fig. S2). *sage* and *fkh* were also both expressed in the venom glands of first instars. However, their signal was weak, making it challenging to distinguish from autofluorescence, especially from the eyes (Fig. 6b).

**Figure 6.**
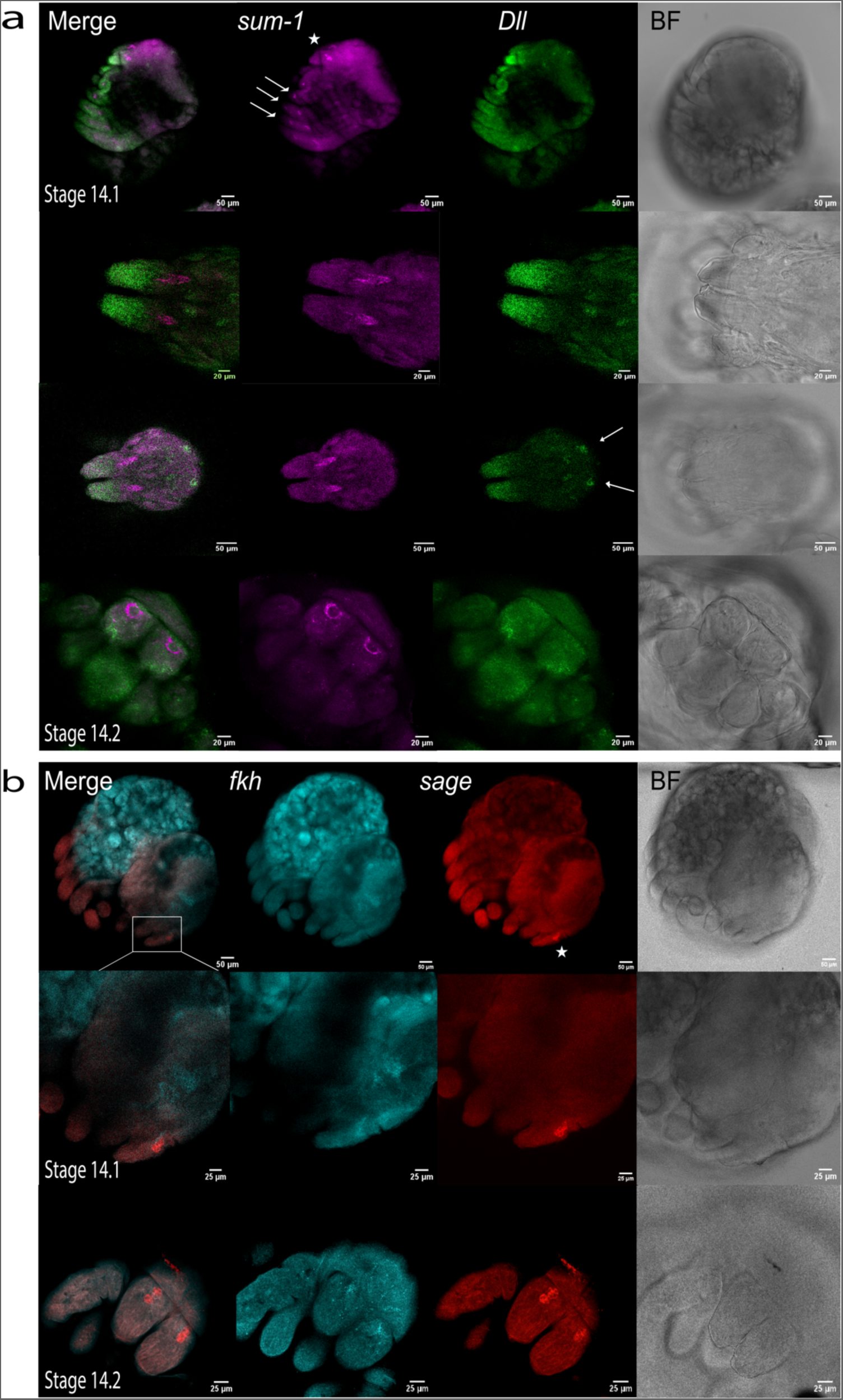
Expression of venom gland markers in the prosoma of stage 14 embryos. a) Expression of *sum-1* and *Dll* in stage 14.1 and 14.2 embryos. Note the additional *sum-1* expression in the leg joints (arrows), and the *Dll* signals in correspondence of the eye primordia (arrows). s*um-1* signal from venom glands marked with a star. Scale bar: 50 *μM*. Scale bar: 20 *μM*. b) HCR signals of *fkh* and *sage* in stage 14.1 and 14.2 embryos. Signal in venom glands marked with a star. BF: brightfield.

In conclusion, progressing from early postembryos to late first instars, we observed a change in marker expression from within the chelicerae towards the dorsal side of the prosoma (Fig. 8).

#### Embryonic stages

Since venom glands were already identified in early postembryos, we extended our investigation to the embryonic stages, starting from the stage just before eclosion and proceeding earlier^20^.

At stage 14, *sum-1* exhibited the previously observed ring-like expression pattern at the proximal end, or base, of the chelicerae (Fig. 6a). We also detected *sum-1* expression in the leg joints (Fig. 6a) and at the level of the coxae (Supplementary Fig. S3). *Dll* was highly expressed at the distal end of the chelicerae, particularly in the fang primordium, as well as in the endites, labium and legs; additionally, we observed signals that appeared to correspond to the eye primordia (Fig. 6a). The toxin gene was not expressed at this stage (Supplementary Fig. S4). Strong *sage* signal was detected within the glands, while *fkh* signal was comparatively weak and not observed in all samples (Fig. 6b). However, *fkh* expression was strong in other regions of the prosoma potentially corresponding to the brain, as well as in the opisthosoma (Fig. 6b).

At stage 13.2, *sum-1* was expressed along a line on the dorsal side of the chelicerae rather than the ring-like pattern observed later (Fig. 7a). Additional expression was observed in the leg joints and coxae. *Dll* was expressed distally in a small region on the dorsal side of the chelicerae (Fig. 7a). In contrast, *Dll* expression in the legs and pedipalps extended for about two-third of their length mainly in the ectodermal cells (Fig. 7a). Notably, punctiform expression of *Dll* resembling *sum-1* were also observed at the level of the coxae, just below *sum-1* signals, although the expression of the two genes did not appear to overlap (Fig. 7a). At this stage, only *sage* appeared to delineate the venom gland rudiments, which were observed distally in the chelicerae (Fig. 7b). Additionally, spotty expression of *sage* was visible in the pedipalps. *fkh* expression was weak and overlapped with *sage* in the gland primordium and was largely expressed in the opisthosoma (Fig. 7b).

**Figure 7.**
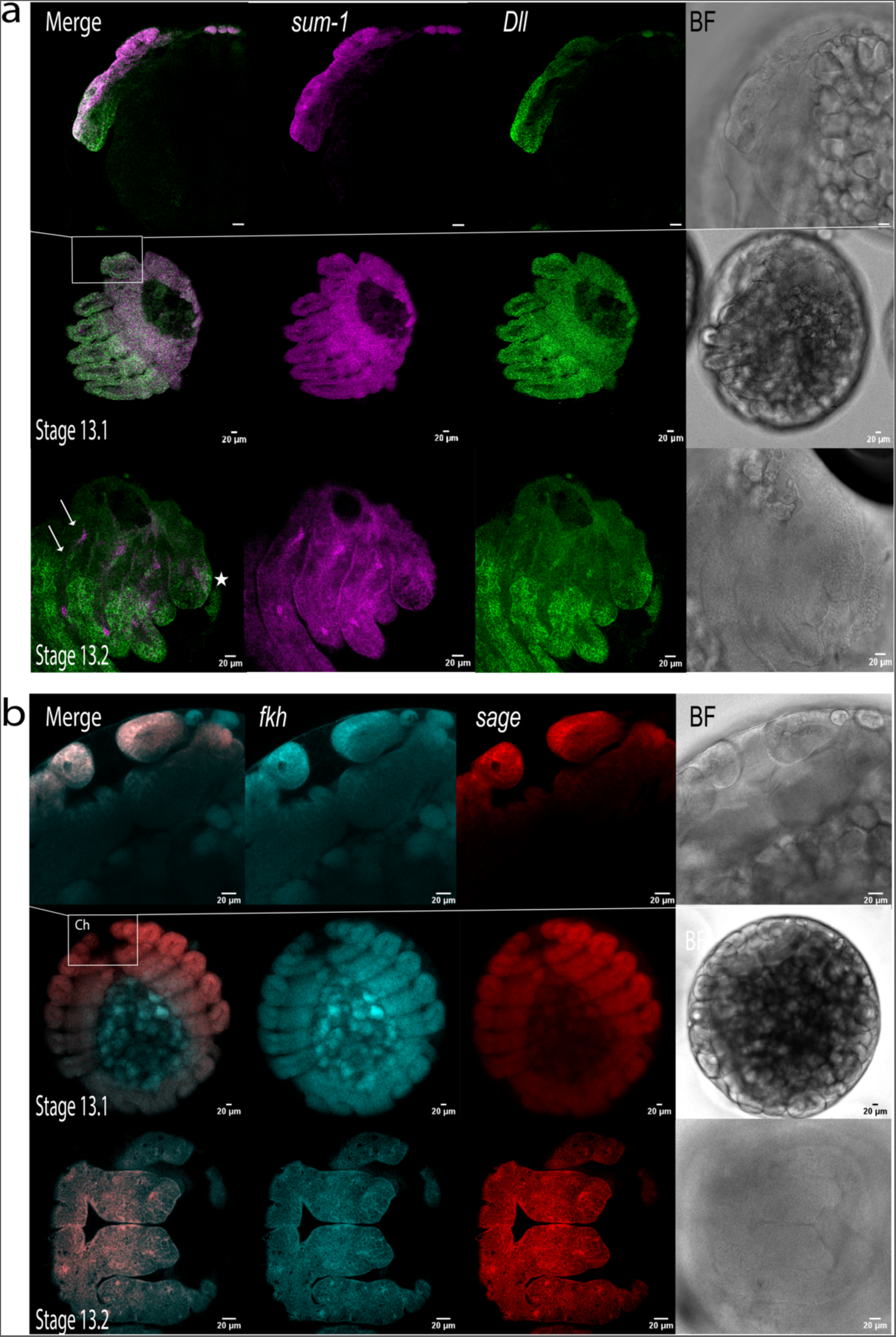
Embryonic expression of venom gland markers in stage 13 embryos. a) Expression of *sum-1* and *Dll* in stage 13.1 and 13.2 embryos. At stage 13.2, additional punctiform signals were observed in the coxae (arrows) along with expression at the dorsal side of the chelicerae (star). Scale bar: 20 μm. b) Expression of of *fkh* and *sage* in stage 13.1 and 13.2 embryos. In stage 13.2, *sage* expression corresponds to the venom gland rudiments, which are now closer to the tip of the chelicerae. BF: brightfield.

At stage 13.1, *sum-1* had a similar expression pattern to stage 13.2, although it seemed more broadly expressed on the dorsal side of the chelicerae (Fig. 7a). *Dll* had a similar expression pattern in the chelicerae (Fig. 7a). *sage* signal did not clearly correspond to the venom glands, but instead it was found expressed along a lumen in the interior of the chelicerae towards the distal end (Fig. 7b). This lumen was also evident when staining with DAPI, and additional brightfield CSLM images revealed the presence of an invagination (Supplementary Fig. S5). *fkh* was also expressed in the chelicerae in a similar pattern to *sage*, although it was more prominently expressed in the opisthosoma as in the subsequent stage (Fig. 7b).

We also assayed the expression of our focal genes at stages 9 and 12, but no venom gland-specific signals were detected (Supplementary Fig. S6,S7).

In summary, the venom gland primordia emerge in embryonic stage 13 at the distal end of the chelicerae, progressing proximally towards the base of the chelicerae by the end of the embryonic development at stage 14 (Fig. 8, Supplementary Fig. S8). Following hatching, the glands continue elongating into the prosoma.

**Figure 8.**
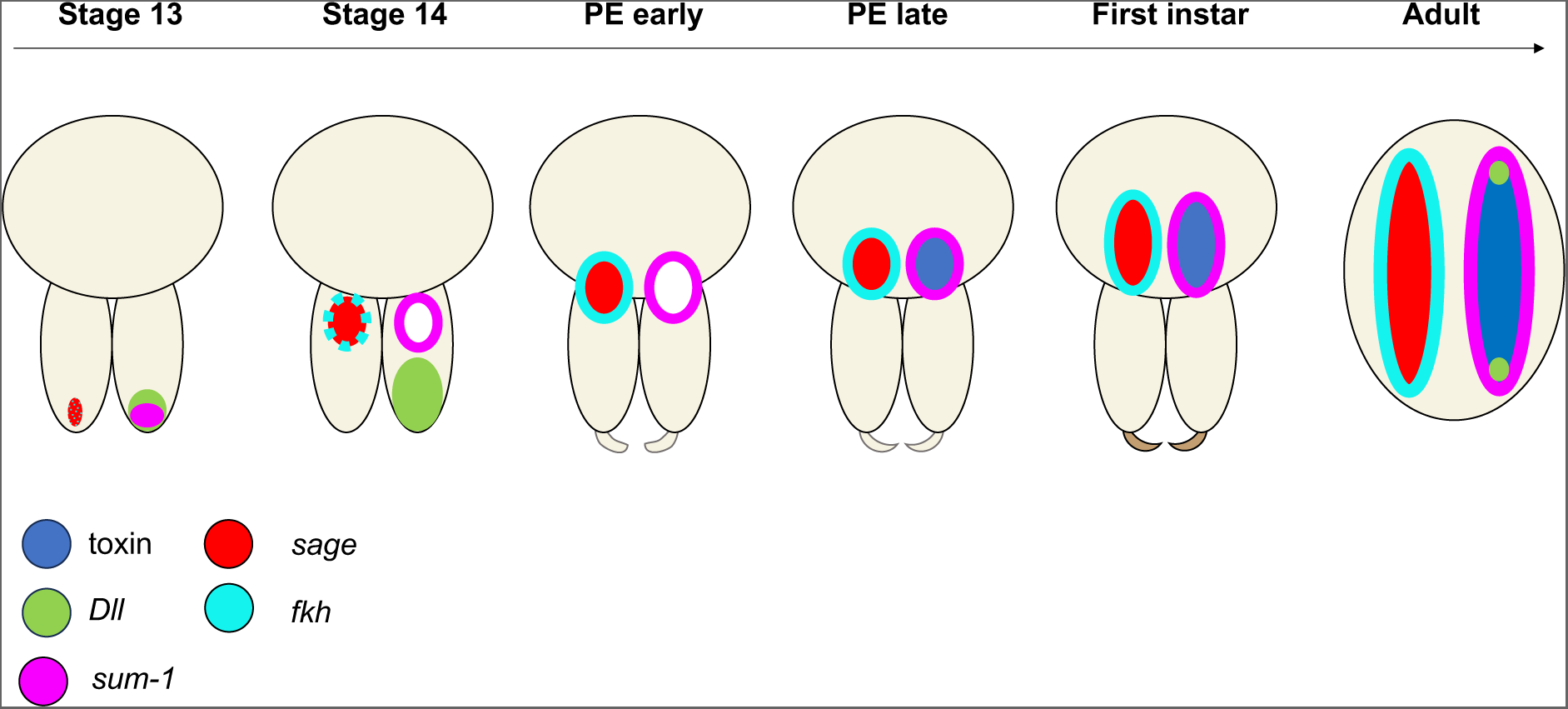
Schematic representation of venom gland marker expression during development. At stages 13 and 14, *fkh* expression is weak and therefore represented as dots. *sage* is exclusively expressed in the gland primordium at stage 13, while *sum-1* has a broader expression at the distal end of the chelicerae. *Dll* is broadly expressed at the distal end of the chelicerae. At stage 14, *sum-1* is specifically expressed in the muscle layer around the gland rudiments. *sage* and *fkh* are also expressed in venom glands. The toxin begins to be expressed in the late postembryos. In postembryos and first instars the expression pattern is similar to the adult, but the venom glands are still close to the chelicerae. P = proximal, D = distal.

## Discussion

Venom systems are one of the most successful adaptations in the animal kingdom and yet, little is known about their evolutionary and developmental origins^6^. In this study, we describe the emergence and development of the venom apparatus in the common house spider.

### Venom apparatus development

We discovered that venom gland primordium is first detectable in the embryo at stage 13 likely as an invagination distally in the chelicerae (Supplementary Fig. S5,S8). This observation aligns with previous findings by Rempel^16^ in *Latrodectus* and conforms to the standard model of organogenesis of exocrine glands, characterised by primitive epithelial ingrowth followed by duct elongation^27^. The gland rudiments gradually migrate proximally towards the base of the chelicerae, where they are located at the time of eclosion. After hatching, the glands continue their elongation into the dorsal side of the prosoma (Fig. 8).

The fangs are not yet apparent in embryos and only become visible in the postembryos. The venom apparatus reaches its final form in the early first instar, coinciding with the spiders leaving the cocoon and starting to build the communal web. At this stage, the fangs are developed, possibly ready to deliver venom (Fig. 2).

Our toxin marker, which represents a major component of the venom secretome of the common house spider^21^ begins to be expressed in the late postembryos, just before the first moult, but not at earlier stages, indicating that the secretory cells are undifferentiated or inactive until this stage. A BLAST search of this and other toxin transcripts against RNA-seq libraries of earlier embryonic stages did not yield any significant hits, indicating that toxins are not expressed in embryos. This finding contrasts with previous reports of toxin expression in eggs of *Latrodectus* species^28–30^. However, these egg toxin transcripts were distinct from the latrotoxins generally found in adult venom, suggesting a distinct function, such as possibly protecting against predators or microbes^29^.

### Patterns of expression of selected marker genes

Our RNA-seq analysis of juvenile venom glands revealed similar expression patterns to the adult venom glands, although we detected a higher number of expressed genes in the juveniles. This is very likely due to the inclusion of the chelicerae during the dissection of the juveniles. Many of the venom gland-specific genes were of unknown function or lacked orthologs in other species. To confidently identify venom glands in this spider, we selected a toxin gene known to be secreted in the venom^21^, and a panel of transcription factors, including *Dll*, *sage*, *fkh* and *sum-1*, which exhibited upregulation in the venom glands.

In embryos, *sum-1* displayed spot-like expression in the leg joints and the coxae. *sum-1* is orthologous to the *Drosophila* gene *nautilus*, and to the mammalian family of bHLH myogenic regulatory factors involved in muscle differentiation^31^. In other venomous animals, the myogenic factors *myf5* or *myod1* were found to be expressed in venom glands, albeit at low levels and not in a tissue-specific manner^22^. This suggests that the specificity of *sum-1* in the muscle fibres surrounding the glandular tissue may be restricted to spiders.

The other marker with a similar expression pattern to *sum-1* was *fkh*, although the expression of this gene was notably lower, particularly in the whole-mount embryos where it was scarcely detected. *fkh* encodes a winged-helix nuclear transcription factor required for salivary gland development and function, and maintains expression of other early-expressed salivary gland transcription factors in *Drosophila*^23,32–35^. Specifically, *fkh*, together with *trachealess* and *hückebein*, regulates the formation of other tubular structures in *Drosophila*, suggesting that these transcription factors function as ‘morphogenic cassettes’ responsible for forming such structures in a variety of tissues^32,34^. Both *trachealess* and *hückebein* orthologs were expressed in the juvenile venom glands, although they were not tissue-specific (Supplementary Dataset S2). Given the tubular shape of spider glands, it is plausible that a similar network has been recruited in spiders.

*fkh* works with *sage* to activate the expression of salivary gland specific gene products, including secreted proteins and their modifying enzymes^23,35^. We found that the spider ortholog of *sage* was exclusively expressed in the venom glands of both juveniles and adults. In particular, *sage* expression was confined to the glandular epithelium across all developmental stages, underscoring the significance of this transcription factor for the development and maintenance of venom glands. *sage* orthologs have been previously found to be upregulated in venom glands of various spider species and scorpions^22^. In insects, *sage* is expressed in robber flies’ thoracic (i.e., venom) glands but not in hymenopteran venom glands^22^, the latter being evolutionary linked to the reproductive system^36^. These observations suggest an evolutionary link between the venom gland and salivary, or any other prosomal gland, that might have been present in the arachnid ancestor. The secretion of toxins similar to those found in spider and scorpion venoms, in the salivary glands of ticks, a parasitic arachnid lineage, may further support this hypothesis^37^.

In addition to its central role in insect salivary gland development, s*age* is also involved in silk gland development in silkworms^38,39^. We examined whether s*age* is expressed in spider silk glands during embryonic stages, but we did not observe any marked signals like in the venom glands (Supplementary Fig. S9). Furthermore, *sage* was not detected in either juvenile or adult silk gland bulk RNA-seq data (Supplementary Dataset S2).

The last marker assayed was *Dll*, which regulates formation of prosomal segments and appendages in *P. tepidariorum*^24–26^. As anticipated, we observed *Dll* expression in the embryonic chelicerae and other appendages. Additionally, we found *Dll* expression in the fangs, and we also detected expression of this gene in adult venom glands, particularly at the extremities. The role of this developmental gene in in this context requires further investigation.

The evolutionary origin of venom glands in spiders remains uncertain and much debated^13,19^. One hypothesis posits that venom glands are modified salivary glands, akin to those of ticks^40,41^. Alternatively, they may have evolved from silk-producing glands present in early chelicerates^13^. Another hypothesis is that venom glands, along with silk glands, could be derived from coxal glands or other prosomal glands, which are quite abundant in other arachnids such as mites^42^.

Zhu and colleagues^19^ advocated the silk gland origin hypothesis based on similarities in the transcriptome profiles of the two organs. However, the absence of key tissues and animal groups in the analysis, such as coxal glands or salivary glands of other arachnid lineages, may have limited consideration of alternative scenarios. The lack of expression of s*age* in the silk gland primordia of *P. tepidariorum*, as well as a lack of expression in adult silk glands in this and other spider species, does not suggest a direct evolutionary link between venom and silk glands. While our markers provide initial insights, spatial expression patterns of additional transcription factors will be crucial to build a more comprehensive understanding of the evolutionary and developmental origins of spider venom glands. Additional expression data from other glands and arachnid lineages will aid in elucidating the evolutionary relationship between these exocrine glands.

## Methods

### Spider husbandry

The *P. tepidariorum* colony was kept at 25°C and humidity of 60% with a gradual light/dark cycle of 16/8h. Adults were kept individually in plastic vials with a coconut husk substrate and fed twice a week with *Musca domestica* flies; the cocoons were transferred into Petri dishes and juveniles fed with *Drosophila sp.* flies.

### Light microscopy

We observed *P. tepidariorum* spiderlings following egg eclosion, specifically focusing on postembryos and first instars, using a Zeiss dissection stereoscope. Specimens were anaesthetised with CO2 prior to dissection of the chelicerae and venom glands.

### Histology

Specimens for histological analysis were fixed in 2.5% glutaraldehyde for 1 hour at room temperature and subsequently fixed overnight at 4°C. After several washes with 0.1 M phosphate buffer, the sample were postfixed in 2% osmium tetroxide solution for 1 hour at room temperature, followed by washing with water, dehydrating through a graded series of acetone, and subsequently a graded series of resin, and embedding using Spurr Low-Viscosity Embedding kit (SIGMA). The samples were incubated in resin blocks at 60°C for 48 hours. Semi-thin sections were obtained with a Leica EM UC7 Ultramicrotome (Leica Microsystem), with a DiATOME diamond knife (Diatome Ltd., Switzerland) at 700 nm thickness. Sections were stained with 0.5% toluidine blue. Images of histological sections were acquired using Zeiss Axio Imager Z2 (Leica Microsystem).

### Bulk RNA-Seq

To identify genes expressed in venom glands suitable as markers for *in situ* HCR experiments, we conducted bulk RNA-Seq. At four days post-hatching (first instars), we dissected the venom glands, the remaining prosoma and opisthosoma from 25 spiders. Given the small size and delicate nature of venom glands, they were left attached to the chelicerae to prevent damage. For the prosoma and opisthosoma, we pooled 5-15 individuals in each biological replicate. For the chelicerae/venom glands, we pooled all the 25 samples and obtained only one replicate. Total RNA was isolated using the TRIzol^TM^ Plus RNA Purification kit (ThermoFisher) following the manufacturer’s instructions, with an additional on-column DNA purification step. Subsequently, seven cDNA libraries were generated using the TruSeq RNA Sample Preparation kit (Illumina) with 150 base pair read length. The libraries were subjected to pair-end sequencing on an Illumina NovaSeq at the Genomic Technologies Facility of the University of Lausanne, Switzerland.

In addition to our samples, we analysed gene expression from published SRA libraries of adult spiders, including venom glands (SRR8755631, SRR8755632), prosoma (SRR8755629, SRR8755630), ovaries (SRR8755633, SRR8755634), and silk glands (SRR8755627, SRR8755628).

Raw reads were assessed with FastQC v0.11.9^43^ and quality filtered with Fastp v0.22.0^44^. Reads shorter than 30 bp were discarded. Reads were then pseudo-aligned to the Ptep_3.0 transcriptome (GCF_000365465.3) using Kallisto v0.48.0^45^ and transcript abundances imported in R v.4.2.2^46^ using the *tximport* package^47^.

Since we had only one replicate for the juvenile venom gland sample, we identified genes expressed in venom glands by selecting those with the highest transcript per million (TPM) value in this sample (Supplementary Dataset S3). Subsequently, we calculated the fold change as the difference in expression between the venom gland and the tissue with the second-highest expression value (calculated as average between replicates). Orthology of *P. tepidariorum* genes with other organisms, i.e., *Drosophila*, was inferred using OrthoDB^48^.

### RNA *in situ* Hybridisation Chain Reaction (HCR)

#### Markers

Based on the bulk RNA-Seq data, we selected four markers encoding a venom toxin and three transcription factors. The toxin (XM_016055901.2) has been identified in the venom proteome of *P. tepidariorum*^21^, indicating its expression at the protein level; the transcription factors were *sum-1* (XM_016072711.1), *forkhead C1-A* (XM_016074417.2), *sage* (XM_043045802.1) and *Dll* (XM_016065828.3). To assess the expression patterns of these markers in other organisms, we examined the orthologs in *Drosophila*, humans and other mammals using resources such as FlyBase^49^, the Human Protein Atlas^50^ and Bgee 15.1^51^.

#### In situ HCR

Specimens were collected at ten different time points using the staging of Mittmann and Wolff^20^, and those defined in this study for the post-eclosion stages. Additionally, we investigated gene expression patterns in dissected adult venom glands. Spiders were anaesthetised using CO_2_ and the opisthosoma removed. Additionally, legs were removed where possible to facilitate the penetration of the fixative and probes. Dissections were performed in 0.1% PBS-Tween-20 (PBS-T). Embryos and post-hatched spiders were dechorionated with 3% bleach for 3-4 min, rinsed with tap water, then rinsed 3 or 4 times with MilliQ water. Subsequently, samples were fixed in 1:1 37% formaldehyde:heptane overnight at 4°C on a rotor at low speed (<20 rpm). The fixative was removed, and ice-cold methanol was poured onto the heptane supernatant and mixed vigorously for 1 min. The methanol/heptane mix was removed and replaced with 100% methanol and sample stored at −20°C.

Before HCR experiments, the samples were rehydrated with a series of graded methanol/PBS-T (75%, 50%, 25%) for 5 min on ice, followed by two washes in PBS-T for 5 min each on ice. Samples were post-fixed in 4% formaldehyde at room temperature for 20 min, and washed four times for 5 min in PBS-T. Dissected venom glands were directly fixed in 4% formaldehyde for 2 hours at 4°C on a rotor at low speed, then washed with PBS-T. At this point, we followed the protocol suggested by Molecular Instruments for generic samples in solution with modifications following Manning & Doe^52^. All the following reagents, including probes, were acquired from Molecular Instruments. Briefly, samples were incubated in 300 μl of pre-warmed probe hybridization buffer for 30 min at 37°C, then hybridised in probe solution at 37°C overnight. Samples were washed four times with pre-warmed probe wash buffer for 15 min each at 37°C. Then, samples were washed four times for 5 min in 5X SSC 0.1% Tween-20 at room temperature. Samples were pre-amplified in amplification buffer for 30 min at room temperature using 4-6 μl of hairpins, which were separately heated at 95°C for 90 sec and cooled to room temperature in the dark for 30 min. Samples were incubated in hairpin solution in the dark at room temperature overnight. Samples were then washed in 5X SSC 0.1% Tween-20, mounted in Vectashield and stored at 4°C on glass slides with coverslips for confocal imaging.

We performed two sets of multi-FISH experiments: set 1 included toxin (Alexa-488), *sum-1* (Alexa-647), and *Dll* (Alexa-594); set 2 included *sage* (Alexa-647) and *fkh* (Alexa-488). For each experiment and developmental stage, we examined up to 15 individuals.

#### Image acquisition and analysis

Z-stack images were acquired with a Stellaris 5 White Light Laser (Leica Microsystems) inverted confocal microscope using 10X, 20x and 40X magnification, and analysed with Fiji (ImageJ2-win64). Three dimensional (3D) reconstructions were performed with Imaris x64 9.8.2.

## Data availability

The RNA-seq libraries have been deposited in the NCBI SRA archive with the accession number PRJNA1087249. The HCR images and the R code to analyse the RNA-seq data have been archived at https://doi.org/10.5281/zenodo.10813380. Additional data are provided as supplementary information files.

## Acknowledgments

This work was supported by a Swiss National Science Foundation COST grant IZCOZ0_205460 to GZ. We thank Anna Schoenauer for initial exploratory dissections; the personnel of the Genomic Technologies Facility for the RNA-seq experiment, the Cellular Imaging Facility for the confocal microscopy, and the Department of Electron Microscopy of the University of Lausanne for their help with the histology preparation.

## Additional information

### Supplementary Information

The online version contains supplementary material available at xxx.

## Author contributions

GZ and APM conceived the study design; GB performed the dissections for RNA extraction; AH perform the HCR and histology experiments, image analysis and signal interpretation; GZ generated the bulk RNA-seq data, perform data analysis, and wrote the manuscript; all the authors contributed to the final version of the manuscript.

