## Supplementary Information for "Venom gland organogenesis in the common house spider"

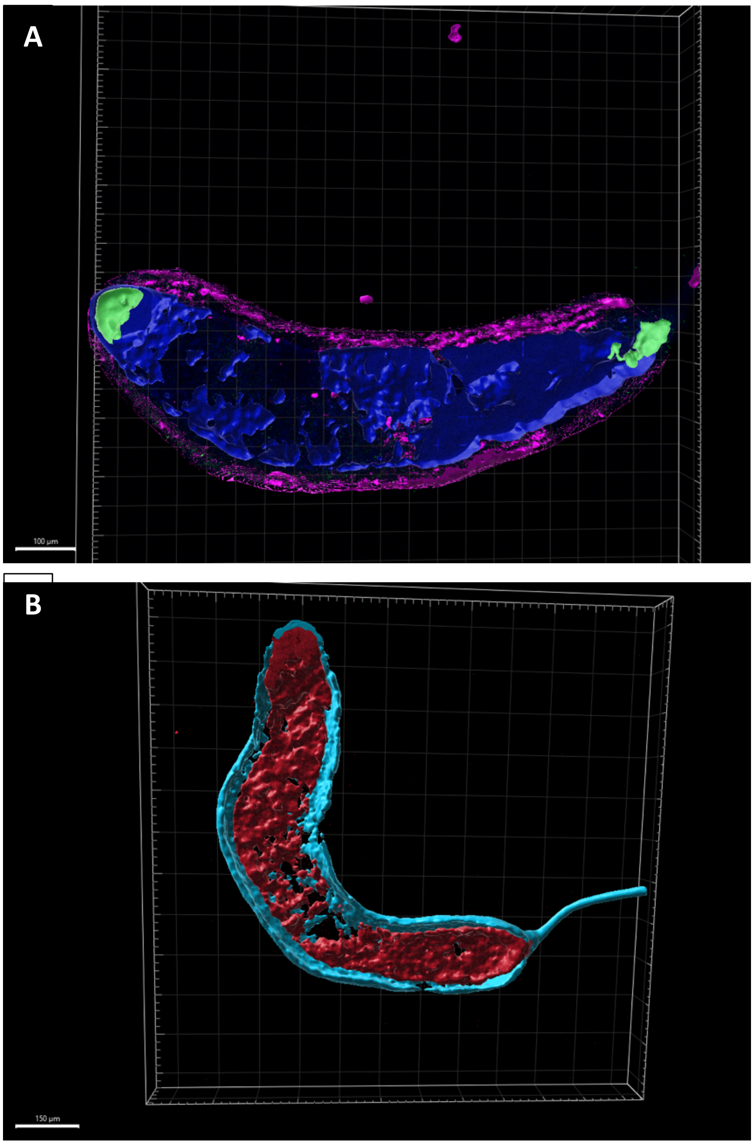


**Figure S1. 3D reconstruction of the *in situ* HCR signals of venom gland markers.** A) Expression of toxin (blue), *Dll* (green), and *sum-1* (pink). Scale bar: 100 *µm*. B) Expression of *fkh* (cyan) and *sage* (red). Scale bar: 150 *µm*.

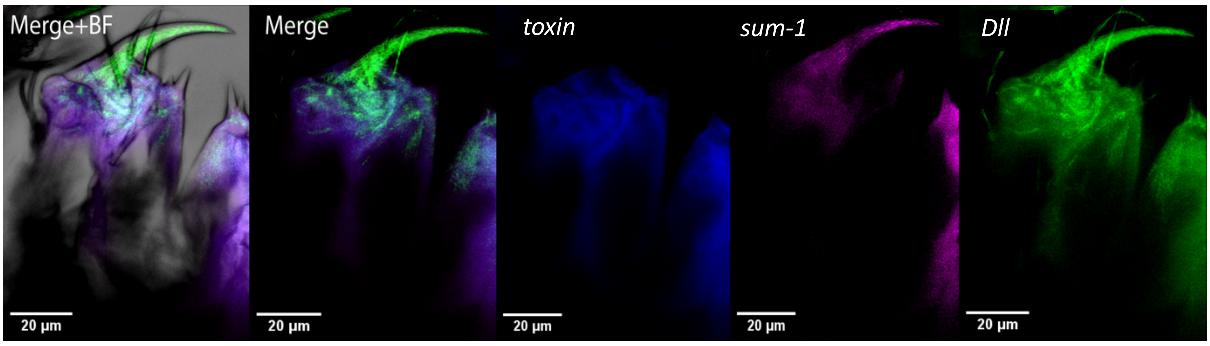


**Figure S2. Expression of the toxin, *sum-1* and *Dll* markers at the tip of the chelicera of whole-mount first instar early stage.** *Dll* is expressed within the fangs, together with *sum-1*, while the toxin gene is not expressed. BF = brightfield.

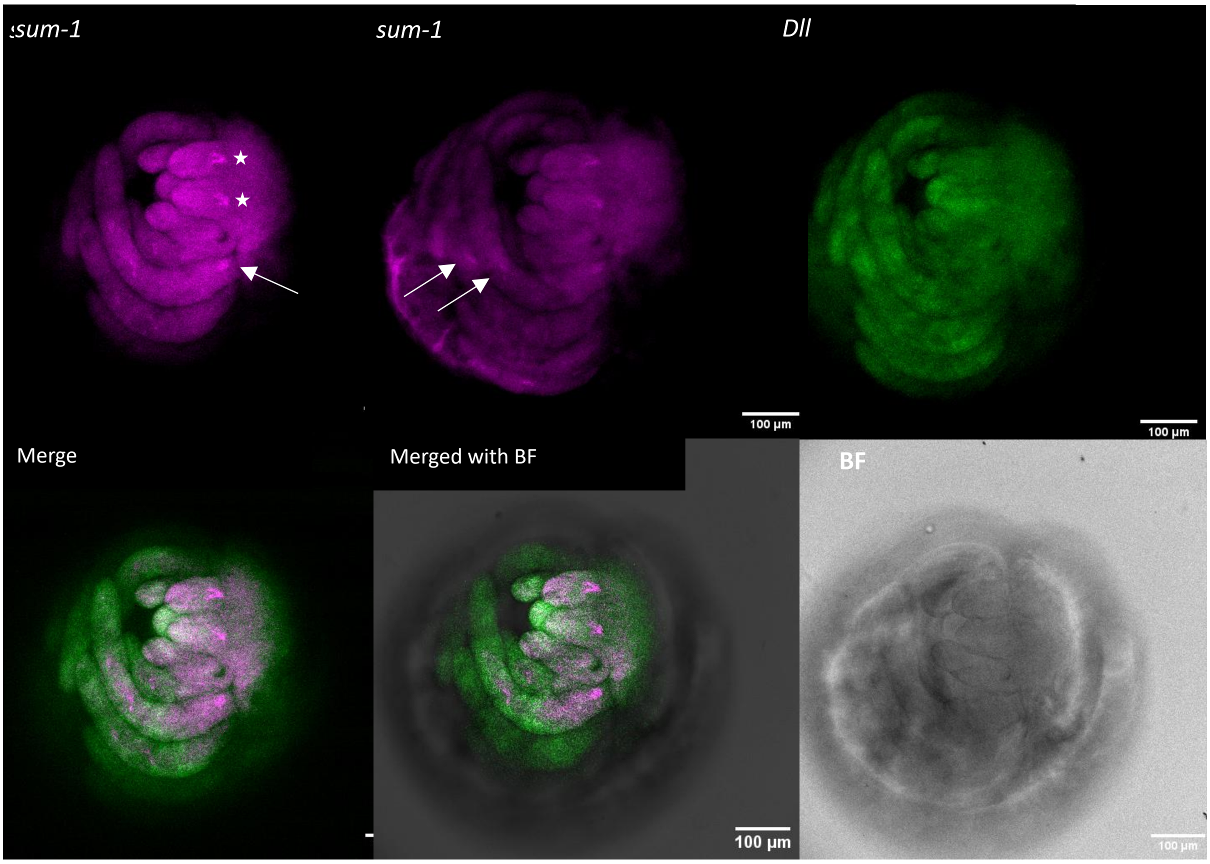


**Figure S3. Expression of *sum-1* and *Dll* at stage 14.** Note the expression of *sum-1* in the coxa of the first leg and the log joints (arrows), along with expression in the venom glands (star) which are located at the base of the chelicerae.

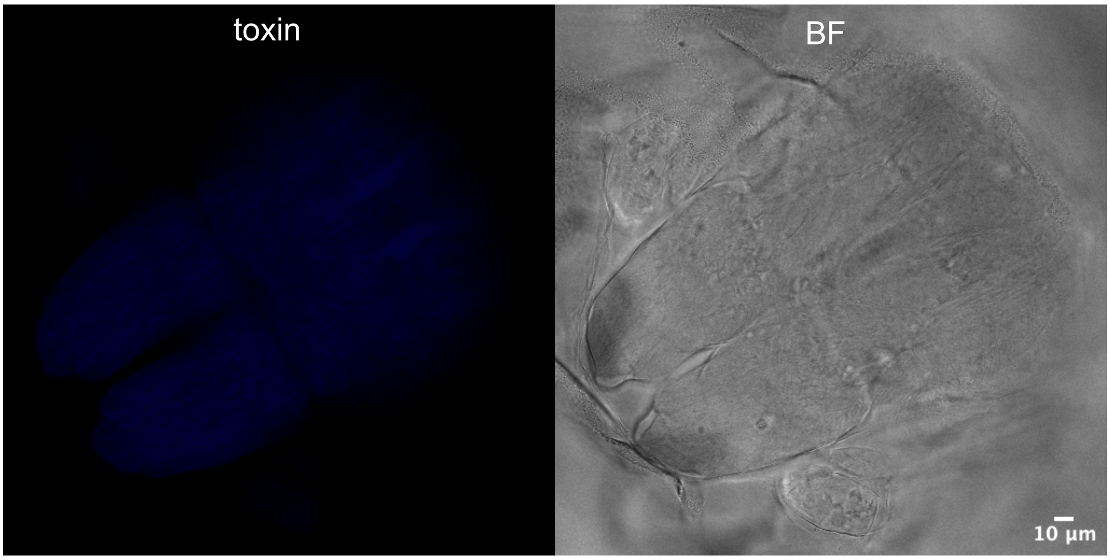


**Figure S4. Expression of the toxin marker in embryos stage 14.** Toxin expression was not detected in the chelicerae nor in the prosoma at this stage. BF = brightfield.

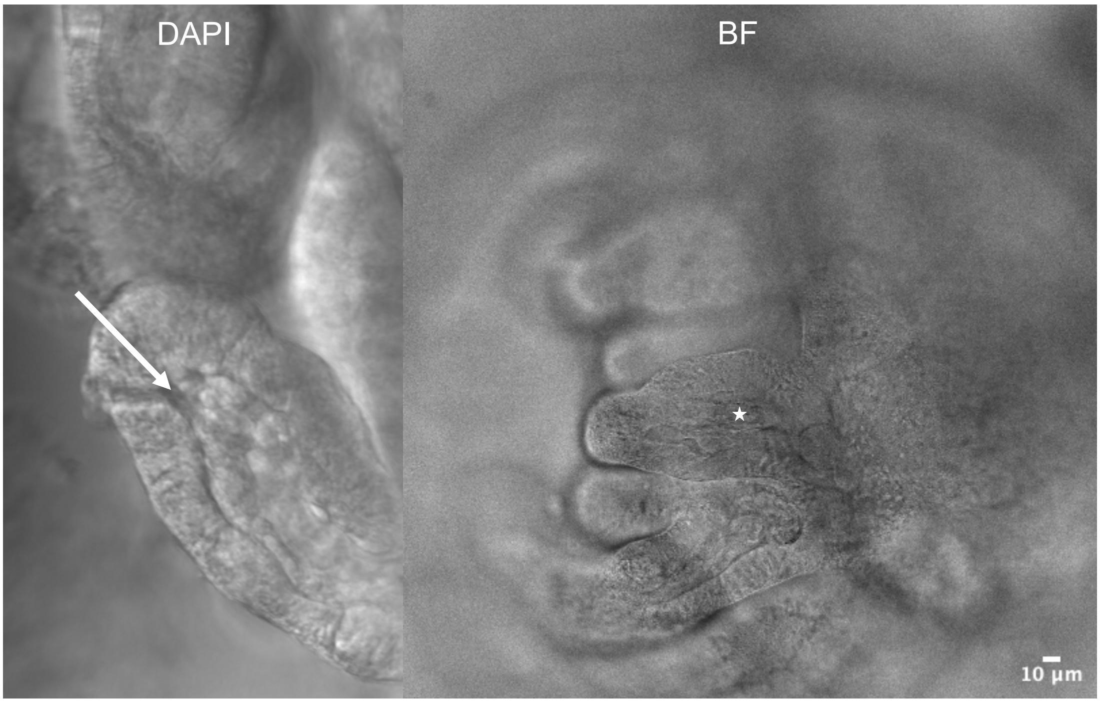


**Figure S5. Interior of the chelicerae of stage 13 embryos.** A) A lumen (arrow) is visible in the distal side of the chelicerae when stained with DAPI. B) An invagination (star) is visible inside the chelicerae in brightfield confocal laser scanning microscopy scan. The images are from two different individuals.

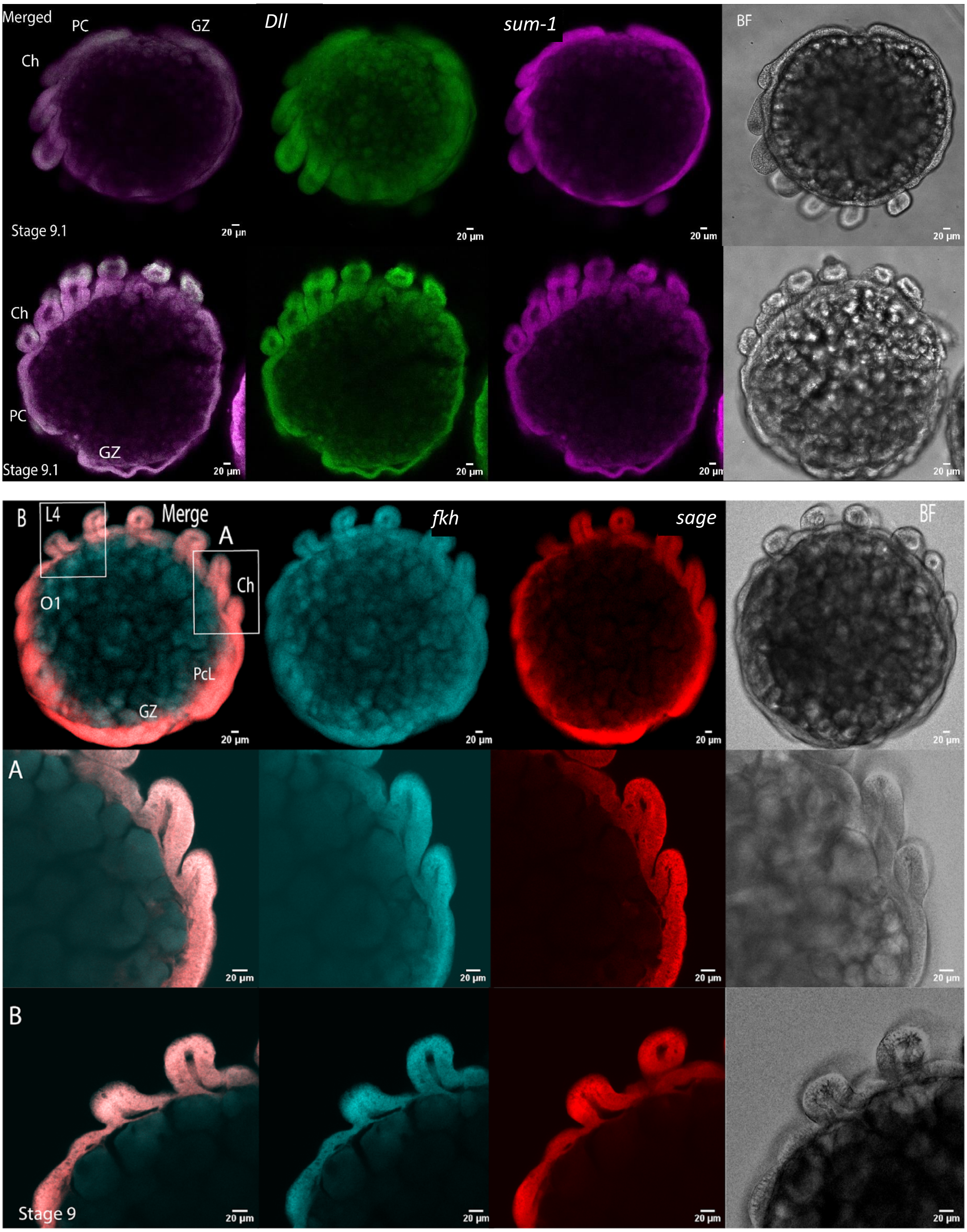


**Figure S6. Expression of marker gene in stage 9 embryos.** The chelicerae are still small buds, and the venom gland primordium has not emerged yet.

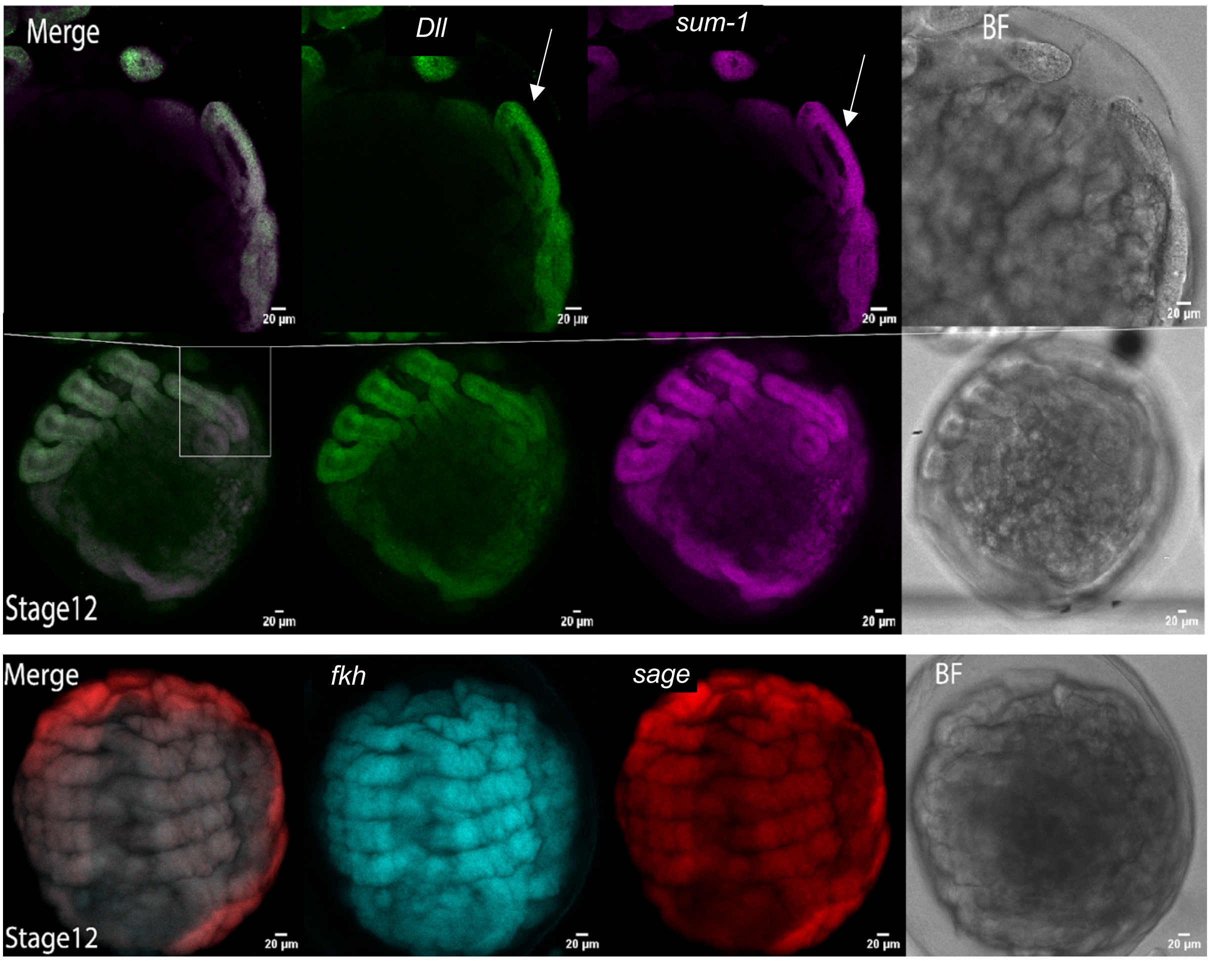


**Figure S7. Expression of marker genes in stage 12 embryos.** At this stage, *Dll* and *sum-1* are expressed on the dorsal side of the chelicerae (arrows). The venom gland primordium has not emerged yet.

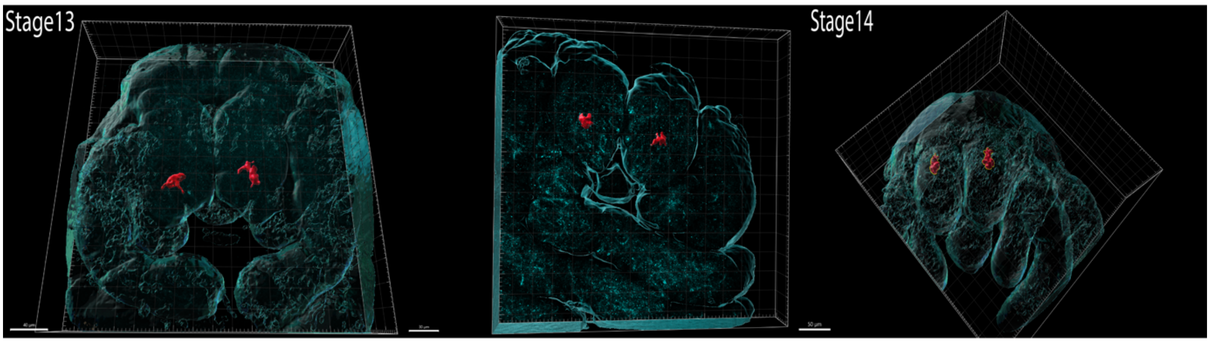


**Figure S8. 3D reconstruction of *sage* expression in embryos.** The venom gland primordium appears at the tip of the chelicerae at stage 13, and during the last embryonic stage it progresses toward the base of the chelicerae.

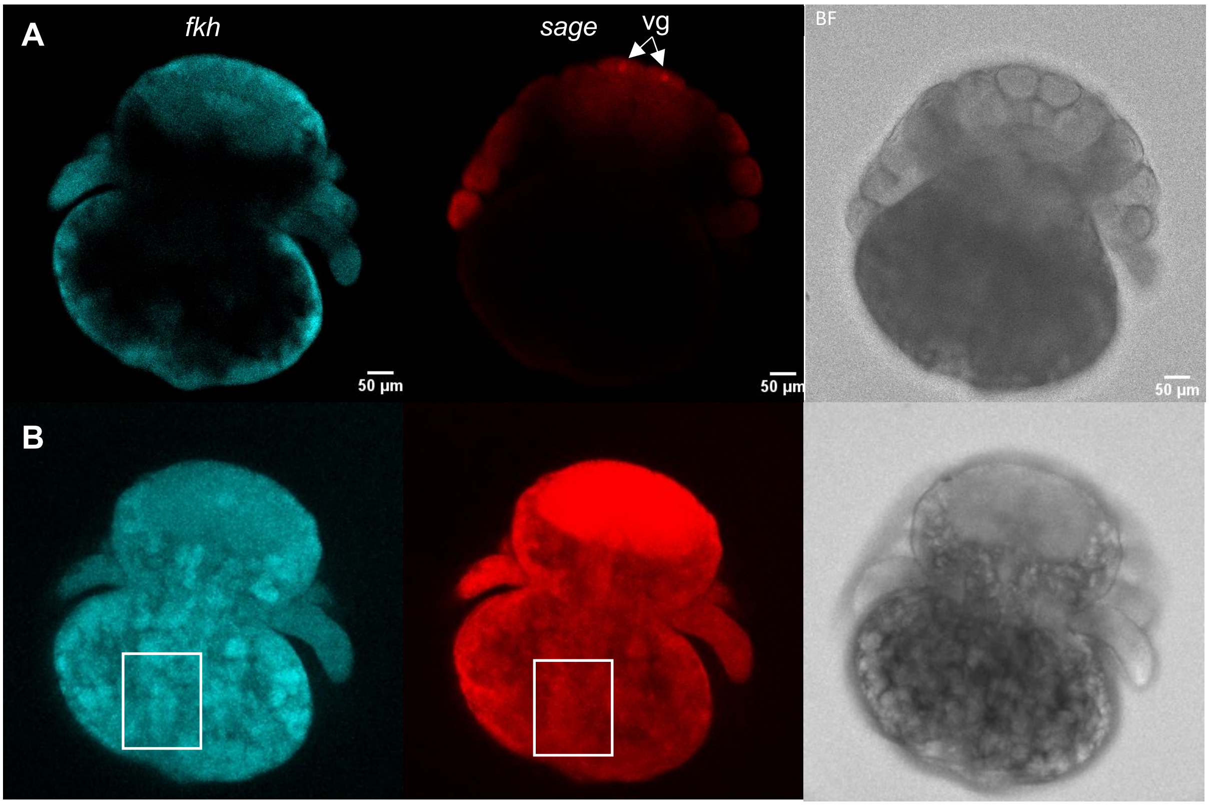


**Figure S9. Expression of *sage* and *fkh* in stage 14 embryos.** A) Focus on the chelicerae to detect expression in the venom glands (vg). B) Focus on the ventral side of the opisthosoma in the area where the silk glands are located (rectangle). No signal corresponding to the silk glands was detected. BF = brightfield.
